# *Klebsiella pneumoniae C*o-infection Leads to Fatal Pneumonia in SARS-CoV-2-infected Mice

**DOI:** 10.1101/2023.07.28.551035

**Authors:** Crystal Villalva, Girish Patil, Sai Narayanan, Debarati Chanda, Roshan Ghimire, Timothy Snider, Akhilesh Ramachandran, Rudragouda Channappanavar, Sunil More

**Author notes:** Corresponding author email: Sunil More, DVM, PhD, DACVP.

## Abstract

SARS-CoV-2 patients have been reported to have high rates of secondary *Klebsiella pneumoniae* infections. *Klebsiella pneumoniae* is a commensal that is typically found in the respiratory and gastrointestinal tracts. However, it can cause severe disease when a person’s immune system is compromised. Despite a high number of *K. pneumoniae* cases reported in SARS-CoV-2 patients, a co-infection animal model evaluating the pathogenesis is not available. We describe a mouse model to study disease pathogenesis of SARS-CoV-2 and *K. pneumoniae* co-infection. BALB/cJ mice were inoculated with mouse-adapted SARS-CoV-2 followed by a challenge with *K. pneumoniae*. Mice were monitored for body weight change, clinical signs, and survival during infection. The bacterial load, viral titers, immune cell accumulation and phenotype, and histopathology were evaluated in the lungs. The co-infected mice showed severe clinical disease and a higher mortality rate within 48 h of *K. pneumoniae* infection. The co-infected mice had significantly elevated bacterial load in the lungs, however, viral loads were similar between co-infected and single-infected mice. Histopathology of co-infected mice showed severe bronchointerstitial pneumonia with copious intralesional bacteria. Flow cytometry analysis showed significantly higher numbers of neutrophils and macrophages in the lungs. Collectively, our results demonstrated that co-infection of SARS-CoV-2 with *K. pneumoniae* causes severe disease with increased mortality in mice.

## Introduction

The COVID-19 pandemic caused by severe acute respiratory syndrome coronavirus 2 (SARS-CoV-2) has affected more than 700 million people with 6.9 million deaths. Most individuals infected with SARS-CoV-2 were asymptomatic or had mild fever, headache, anosmia, fatigue and mild respiratory symptoms [1–3]; however, a significant number of patients exhibited severe disease and required hospitalization. The hospitalized patients showed pneumonitis, hypoxia, acute respiratory distress syndrome (ARDS), and multiple organ failure [4, 5].

Viral respiratory tract infections such as influenza and SARS-CoV-2 are commonly accompanied by co- or secondary infections with other viruses, bacteria, and/or fungal pathogens [6, 7]. Bacterial coinfections were responsible for high mortality during the 1918 influenza pandemic [8]. Patients with severe respiratory signs are at a higher risk of secondary infections due to compromised respiratory epithelial barrier or dysregulated immune system [9, 10]. Virus infections induce upregulation of pro-inflammatory cytokines, which can lead to epithelial cell damage or death [11], resulting in the breakage of the physical epithelial barrier and increase bacterial attachment and colonization [12]. During the COVID-19 pandemic, high numbers of SARS-CoV-2 patients had detectable bacterial co-infection, and the most common bacterial agents identified were *Staphylococcus aureus*, *Streptococcus pneumoniae*, and *Klebsiella pneumoniae* (*Kp*) [13, 14]. Additionally, secondary infections and coinfections can also contribute to acute respiratory distress syndrome (ARDS) [15] and leading up to 80.5% fatality in these patients [16].

*Kp* causes hospital-acquired infections and is the third most common gram negative infection [17]. Clinical studies have also identified *Kp* infection in COVID-19 patients with secondary bacterial pneumonia and mortality [13, 18, 19]. *Kp* was detected in 37 to 55 % of COVID-19 patients, and the highest incidence was noted during early days of SARS-CoV-2 infection [13, 20]. Despite the high rate of co-infection, no study has evaluated the effect of *Kp* co-infection on COVID-19 pathogenesis and clinical outcome. Murine models for SARS-CoV-2 have been established that exhibit similar symptoms and immune responses to human patients and are valuable tools for studying pathogenesis [21, 22]. These models have been used to study SARS-CoV-2 co-infections, demonstrating the importance of secondary bacterial infections that can lead to high lethality during SARS-CoV-2 infection [23, 24].

In this study, we employed a mouse model to investigate the impact of SARS-CoV-2 and *Kp* co-infection. Our findings revealed a significantly higher mortality rate among co-infected mice compared to those infected with SARS-CoV-2 or *Kp* alone. Moreover, we observed a marked escalation in lung pathology, bacterial load, and infiltration of immune cells in co-infected mice. This mouse model can serve as a crucial tool for unraveling the intricate pathogenesis underlying both SARS-CoV-2 infection and secondary bacterial infections.

## Material and methods

### Respiratory pathogens PCR array

Deidentified and leftover SARS-CoV-2 positive nasopharyngeal swabs (n = 50) collected from human patients (IRB-20-357-STW) received at the Oklahoma Animal Disease Diagnostic Laboratory (OADDL) for COVID-19 testing between April 2020 – July 2020 were used in this study. Total RNA was extracted from these patient samples using the KingFisher Flex platform (Thermo Fisher Scientific, MA) and a commercially available kit (MagMax viral/pathogen nucleic acid isolation kit; Thermo Fisher Scientific) following the FDA Emergency Use Authorization protocol provided by the manufacturer. A commercially available multiplex real-time PCR-based kit (Fast Track Diagnostics, Sliema, Malta) was used for detecting other respiratory pathogens. This PCR can detect 32 of the most common respiratory pathogens, including viral, bacterial, and fungal agents. The list of pathogens includes influenza A virus; influenza B virus; influenza C virus; influenza A (H1N1) virus (swine-lineage); human parainfluenza viruses 1, 2, 3, and 4; human coronaviruses NL63, 229E, OC43, and HKU1; human metapneumoviruses A/B; human rhinovirus; human respiratory syncytial viruses A/B (HRSV); human adenovirus; enterovirus; human parechovirus; human bocavirus; *Pneumocystis jirovecii; Mycoplasma pneumoniae; Chlamydophila pneumoniae; Streptococcus pneumoniae; Haemophilus influenzae B; Staphylococcus aureus; Moraxella catarrhalis; Bordetella spp.; Klebsiella pneumoniae; Legionella pneumophila/longbeachae; Salmonella* spp. The PCR results were interpreted as detected or non-detected.

### Bacteria

*K. pneumoniae* subspecies *pneumoniae* (Schroter) Trevisan (Cat #8045 ATCC, Manassas, VA) was used for mice infection studies. Bacteria were grown in nutrient broth (BD, Franklin Lakes, NJ) to an exponential growth curve (OD600 = 0.6). The bacterial stock was diluted in phosphate-buffered saline (PBS) pH 7.4 (Gibco, Evansville, IN) and prepared to a final range of 100-200 colony-forming units (CFU) per mouse (in 50 µl) and transported on ice until inoculation. After mice infection, 50 μL of remaining stock was plated on nutrient agar (NA) (BD, Franklin Lakes, NJ) and grown at 37 °C overnight to obtain an estimate of bacteria delivered to each mouse.

### Virus

Mouse-adapted (MA) SARS-CoV-2 strain obtained from Dr. Stanley Perlman (University of Iowa) was used in this study. The virus was propagated in Vero E6 TMPRSS2 T2A ACE2 cells (Cat #NR-54970 BEI, Manassas, VA) and maintained in Dulbecco’s Modified Eagle’s Medium supplemented with 10% fetal bovine serum (GIBCO, Evansville, IN) and 10 µg per mL puromycin (InviovGen, San Diego, CA) at 37 °C in a humidified 5% CO_2_ incubator. The infected cells were clarified by centrifuging at 2000 rpm for 5 minutes. Serial dilutions were prepared from supernatants and 100 µl was added to the Vero E6 cell (ATCC, CRL-1586) monolayers for 1 hour with gentle agitation every 10 minutes. After an hour of incubation, the media was removed and replaced with an overlay composed of Avicel (Du Pont), 2XDMEM (Millipore Corp., Burlington, MA), and DMEM with a 10% FBS (Gibco, Evansville, IN). Cells were incubated at 37 °C in a humidified 5% CO_2_ incubator for 3 days. The following incubation, overlay was removed, and cells were fixed with 10% formalin for 5 minutes and stained with 0.05% crystal violet to visualize plaques. All tests were performed in duplicate. The viral load in mouse lungs was estimated using a similar plaque assay. The left lung lobe was homogenized in 500 μL of Opti-MEM using a bead mill and centrifuged at 2000 rpm for 5 minutes. Ten-fold serial dilutions were used for plaque assay. To prevent the growth of *Kp* in the Vero E6 cells during plaque assay, 40 μg/mL of Gentamicin (GIBCO, Evansville, IN) was used during virus incubation and in overlay media.

### Mice studies

Eight to 9-weeks-old female BALB/cJ mice were purchased from Jackson Laboratories (Jackson Research Laboratories, Bar Harbor, ME). Mice were housed in biosafety cages in an animal BSL-3 facility at Oklahoma State University. All the experiments were approved by the Oklahoma State University Institutional Animal Care and Use Committee. Mice were divided into the following groups: control (PBS), S-CoV-2 (S-CoV-2), *K. pneumoniae* (*Kp*), and S-CoV-2+ *K. pneumoniae* (S-CoV-2 + *Kp*). On day 0 under isoflurane anesthesia, the S-CoV-2 and S-CoV-2 + *Kp* groups were intranasally inoculated with 250–1000 PFU of MA-SARSCoV-2, whereas the control and *Kp* groups were administered 50 μL PBS. Days post infection (dpi) were based on SARS-CoV-2 inoculation day. On 4 dpi, *the Kp* and S-CoV-2 + *Kp* groups were inoculated intranasally with 100–135 CFU of *Kp* under isoflurane anesthesia. Mice were weighed and scored daily for clinical signs for 12 dpi. Clinical score parameters were: 1 = normal skin and active in the cage before handling; 2 = ruffled skin and alert; 3 = hunched posture, skin tent and decreased resistance to handling; 4 = piloerection with severe skin tent, only moves when touched; 5 = failure to right itself, nonresponsive. Mice were humanely euthanized if they lost more than 25% of their starting weight or if they had a clinical score of 4 or more. Survival was defined as an animal not losing 25% of the starting body weight, sick, or dead. Animals assigned a day for sacrifice were included in the count if they fit the criteria. All other animals that did not fit the criteria during the assigned sacrifice were excluded from the numbers. Necropsies were performed on sick mice and mice that were assigned for sample collection. The lung lobes were collected for histopathology, flow cytometry, and bacterial, and viral load analysis.

### Determination of bacterial load in lungs and heart blood

Approximately 10 μL of heart blood was collected aseptically and plated directly on NA plates during necropsy. The plates were incubated at 37 °C and 5% CO_2_ overnight. The left lung was homogenized using a bead mill (Fisher, Hampton, NH) in 500 μL of Opti-MEM (Gibco, Evansville, IN). Ten microliters (in triplicate) of 10-fold serially diluted lung homogenates were plated on NA plates. The plates were incubated overnight at 37 °C and 5% CO_2_ and CFUs were counted.

### Flow cytometry

The phenotypic profile of lung-infiltrating immune cells was analyzed in the left lung lobe. For this, lungs were treated with collagenase-D and DNAse1, and isolated cells were surface-immunolabelled for AM (CD45+ CD11b-CD11c+ SiglecF+), neutrophil (CD45+ CD11b+ Ly6Ghi), inflammatory monocyte (CD45+ CD11b+ Ly6chi), dendritic cell (CD45+ CD11c+ MHCII+), natural killer cell (CD45+ CD3−ve NKP46+), and T-cell markers, and analyzed by flow cytometry. For cell surface staining, lung cells were labeled with the following fluorochrome-conjugated monoclonal antibodies: PECy7 α-CD45 (clone: 30-F11); FITC α-Ly6G (clone: 1A8, BD Biosciences); PE/PerCp-Cy5.5 α-Ly6C (clone: AL-21 [BD Biosciences or clone: HK1.4); V450 α-CD11b (clone: M1/70); APC α-F4/80 (clone: BM8) (unless otherwise stated, all from eBioscience). The labeling of the cell surface and intracellular markers was performed as previously [25]. All fluorochrome-conjugated antibodies were used at a final concentration of 1:200 (antibody: the FACS buffer), except for FITC-labeled antibodies used at 1:100 concentration.

### Histopathology

The lung lobes were perfused with 200 µl of 10% formalin and stored for 72 hours to inactivate the virus. Lungs were then trimmed and processed for hematoxylin and eosin (H&E) staining. Histopathological lesions were scored by two American College of Veterinary Pathology Board-certified pathologists in a blinded fashion. The lesions scored included bronchiolitis, thrombosis, fibrin, necrosis, interstitial pneumonia, edema, and hemorrhages. The lesions were scored on a scale of 0-4; 0 = no lesion, 1 = minimum, 2 = mild, 3 = moderate, and 4 = severe. The sum of all lesion scores for each mouse was used for data plotting and analysis.

### Data Analysis

All statistical tests were performed on GraphPad Prism 9 (San Diego, CA) using a one-tail t-test, Welch’s test, and Kaplan–Meier survival curve with a Mantel-Cox test.

## Results

### *Klebsiella pneumoniae* is the most common coinfecting pathogen in COVID-19 patients

We identified seven different pathogens in 54% (27 of a total 50) of the samples tested. The remaining 23 samples were negative for the pathogens included in the assay. The seven pathogens were: Human Respiratory Syncytial Virus (HRSV) A and B, *Staphylococcus aureus* (*Sa*), *Klebsiella pneumoniae*, Enterovirus, *Pneumocystis jirovecii*, *Salmonella sp*., and *Moraxella catarrhalis* (Fig. 1A). Out of the 27 samples, 26 samples contained at least one bacterial agent, 2 samples contained two viruses (HRSV and Enterovirus), and 1 sample contained a fungal agent (*P. jirovecii*) (Fig. 1B). Of the SARS-CoV-2 positive samples (27 samples), fifteen samples contained a single infectious agent, 10 samples contained two pathogens, and 1 sample contained three pathogens (Fig. 2C). The most common bacterial agents observed were *Kp* (38%) and *Sa* (28%) (Fig. 1A). Additionally, we found that *Kp* was detected in 18% of samples as a potential standalone infection. Due to its implications in SARS-CoV-2 patients as a secondary bacterial infection [18, 19, 26], we focused on *Kp* for further studies.

**Figure 1.**
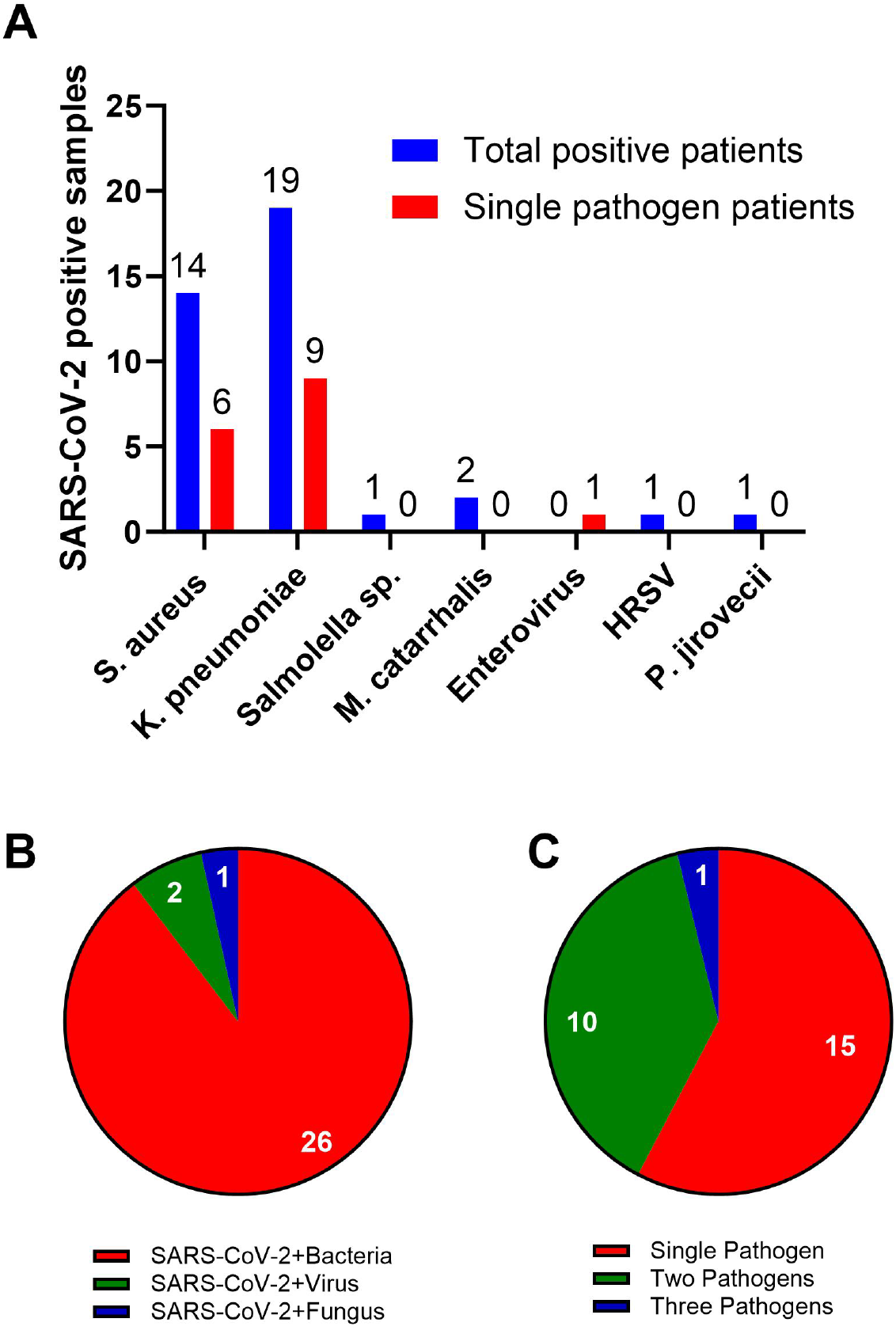
Most common co-infections in SARS-CoV-2 patients. (A) FTD-33 Real-time PCR assay demonstrating pathogens that are detected in SARS-CoV-2 positive nasopharyngeal swabs. Each bar represents number of patients samples; total positive patients (blue bar) indicates presence of pathogen listed on X-axis along with SARS-CoV-2 and other pathogens included in the assay and Single pathogen patients (red bar) indicates presence of pathogens listed on X-axis and SARS-CoV-2. Pie charts demonstrating distribution of co-infecting pathogen type (B) and multiples of infection (C) along with SARS-CoV-2.

**Figure 2.**
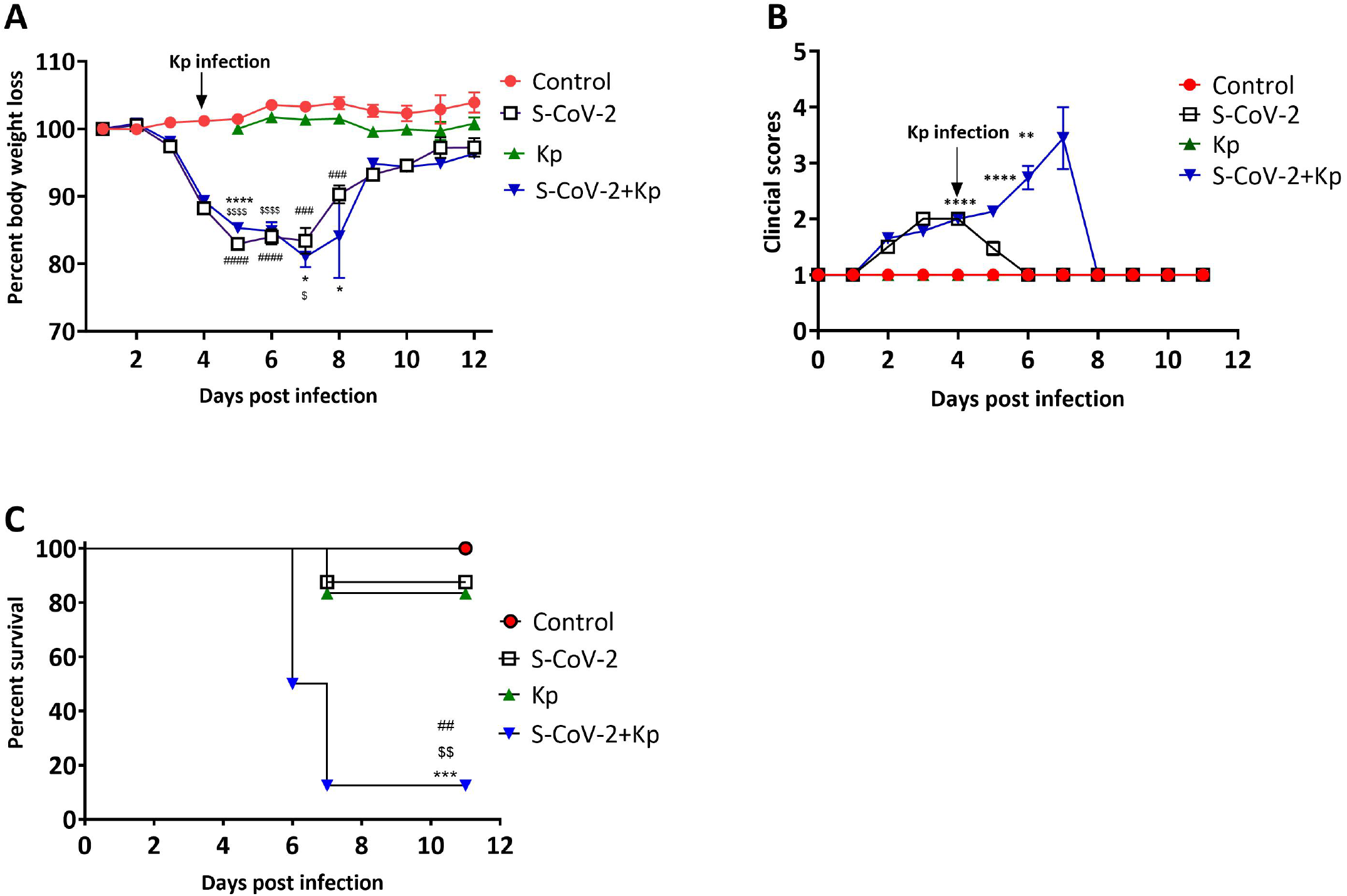
Co-infection with SARS-CoV-2 and *Kp* increases disease severity. (A) Percent body weight loss calculated based on day 0 body weight is presented on y-axis. The data is represented as mean ± SEM. (B) Clinical scores for each group are represented as mean ± SEM (C) Survival of the animals was monitored for 12 days post infection. Data represent n = 18-31 mice per group with two-three independent experiments. Statistical significance was determined by (A) students t-test with *, *P* < 0.05 for S-CoV-2 + *Kp* vs. *Kp*.; ###, *P* < 0.005 and ####, *P* < 0.0005 for Control vs. S-CoV-2 and $$$$, *P* < 0.0005 for Control vs. S-CoV-2 + *Kp*. (B) **, P < 0.005 and ***, P < 0.0005 for S-CoV-2 vs. S-CoV-2 + *Kp*. (C) P < 0.032 based on Kaplan-Meier survival curve with a Mantel-Cox test; *** for S-CoV-2 vs. S-CoV-2 + *Kp; ## for Kp* vs. S-CoV-2 + *Kp and $$* for Control vs. S-CoV-2 + *Kp*.

### Co-infected mice exhibit high rates of morbidity and mortality

We observed a similar pattern of weight loss until day 6 post-infection between the S-CoV-2 and S-CoV-2 + *Kp* groups (Fig. 2A), with both groups of mice losing up to 20% of their initial body weight. However, the S-CoV-2 + *Kp* group, 87.5% of mice continued to lose body weight and succumbed to infection by day 7 (Fig 2C). In comparison, the S-CoV-2-only and *Kp*-*alone* group had only 12.5% and 16.67% of mice died, respectively, due to infection. The difference in outcome can be observed in Figure 2B, which shows a continued increase in clinical score for co-infected mice compared to S-CoV-2-only infection. These findings demonstrate that SARS-CoV-2 co-infection with *Kp* significantly increases clinical signs, disease severity, and mortality.

### SARS-CoV-2 infection promotes *Kp* growth in the lungs

Given the high morbidity and mortality in the co-infection group, we investigated the cause of this phenotype. We determined bacterial load in the lungs of single-(*Kp*) and co-infected mice (S-CoV-2 + *Kp*) and viral load in the lungs of S-CoV-2-only as well as co-infected mice (S-CoV-2 + *Kp*) on 6^th^ dpi. The day 6 post S-CoV-2 was selected because of the significant percentage of death in co-infected mice (Fig. 2C) on that day. We also wanted to examine whether these groups of mice had *Kp* in the bloodstream indicative of septicemia. The number of mice with *Kp* in their heart blood was higher in S-CoV-2 + *Kp* group (*Kp* =1/8 vs S-CoV-2 + *Kp* =7/8). A comparison of bacterial load in the lungs of *Kp* and S-CoV-2 + *Kp* showed a significant difference (*p* = 0.0343) with *Kp* mice having mostly no bacterial growth in the lung (Fig. 3A). Virus titer in the lungs of S-CoV-2 and S-CoV-2+ *Kp* groups, was did not show significant difference (Fig. 3B). Thus, the high mortality rate in the S-CoV-2 + *Kp* groups are associated with increased *Kp* replication rather than increased SARS-CoV-2 replication.

**Figure 3.**
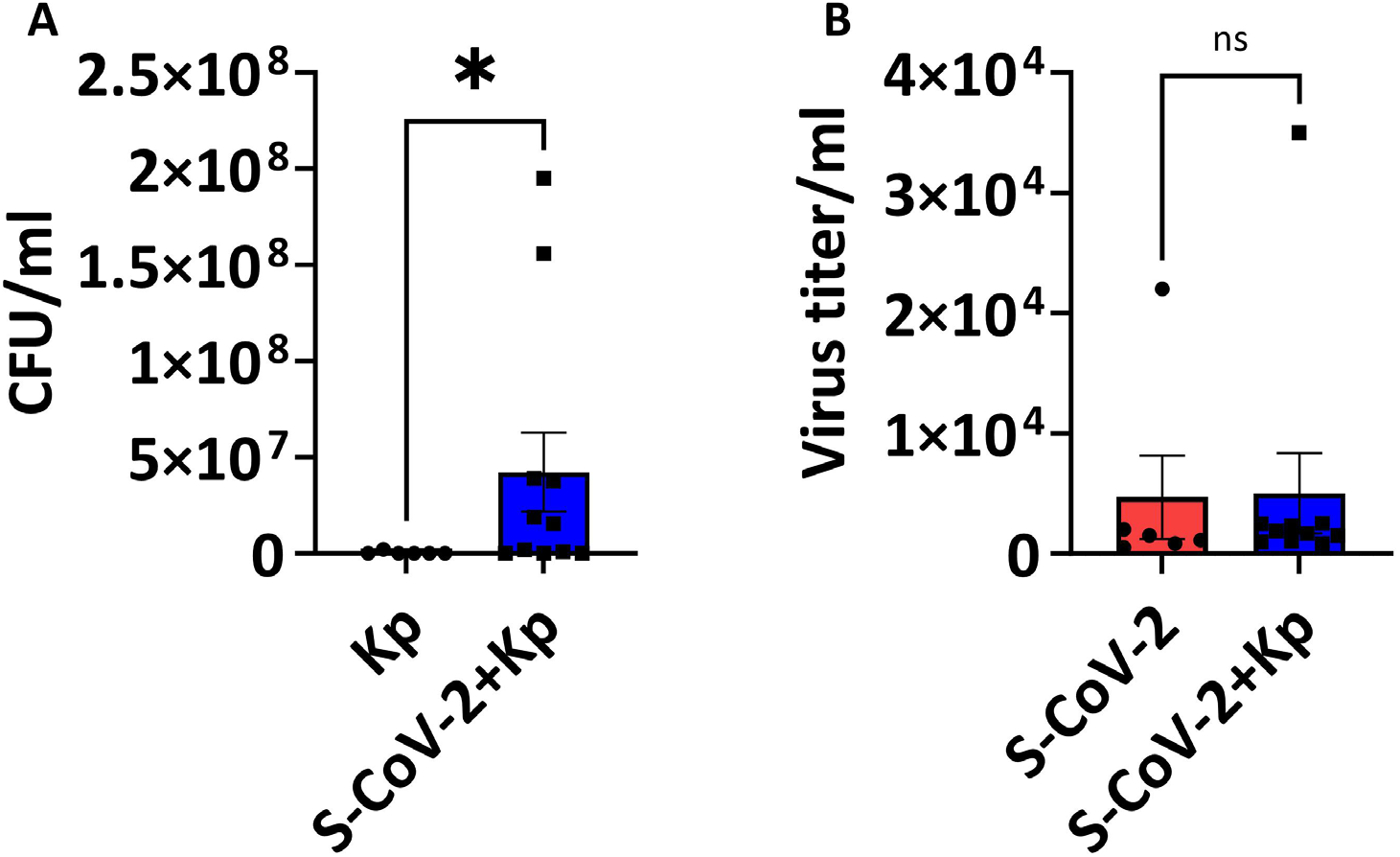
SARS-CoV-2 co-infected mice exhibit increased *Kp* propagation in the lungs. SARS-CoV-2 infected mice were coinfected with *Kp* or *Kp*-alone on day 4 and lungs were collected on day 6 to examine the viral and bacterial loads. Bacterial count (A) and viral titers (B) in the lungs of respected group are presented. Data is presented as mean ± SEM of 2 independent experiments (n = 6-11). Statistical significance was obtained with Welch’s test with * P < 0.05.

### Co-infection causes excessive infiltration of neutrophils and macrophages into the lungs

Since COVID-19 patients with ARDS contain a higher percentage of neutrophils [27] as well as their phagocytic role in bacterial infection, we wanted to quantify the neutrophil response in the lungs. Neutrophil recruitment in the lungs was significantly higher in the S-CoV-2 + *Kp* co-infection group compared to the other groups (Fig. 4A-C). Furthermore, we wanted to quantify inflammatory macrophages/monocyte changes in the lungs due to infection because of their pathogenic role in mice infected with a lethal dose of SARS-CoV and MERS-CoV [28, 29]. The percentage of inflammatory macrophages/monocytes (IMMs) was not significantly different between all the groups (Fig. 4D-E), however, there were significantly higher numbers of IMMs in the S-CoV-2 + *Kp* group compared to all other groups (Fig. 4F). These increases in inflammatory cells indicate a response to the infection, however, it appears that co-infected mice are recruiting more cells due to the secondary infection, which could be causing immunopathology to epithelial cells.

**Figure 4:**
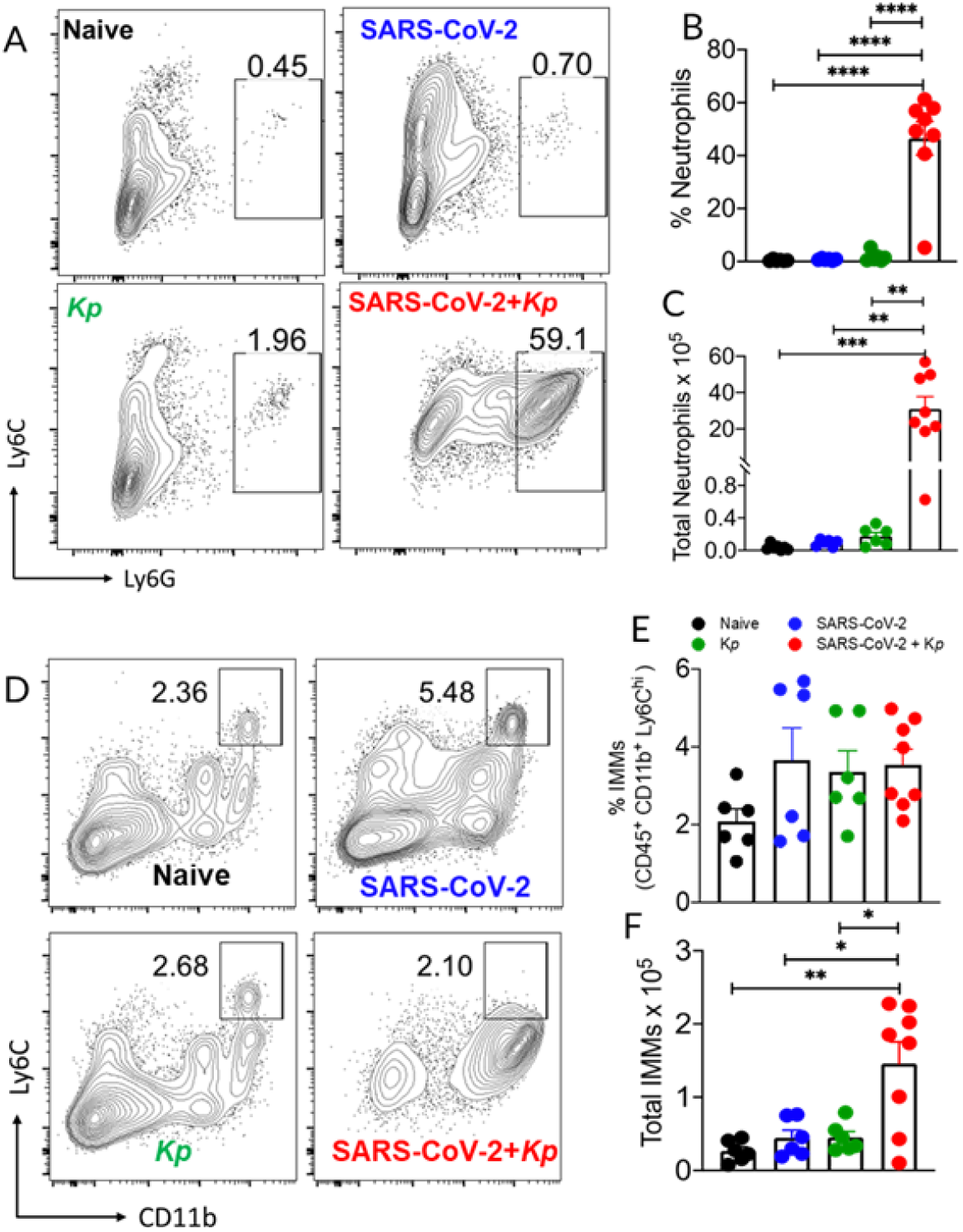
Neutrophil and inflammatory monocytes and macrophages (IMM) responses to coinfection. Representative FACS plots (A and D), quantification of percent (B) and total neutrophils (C), percent CD11b+Ly6Chi IMMs (E) and total CD11b+Ly6Chi IMMs (F) in the lungs. Data were pooled from 2 independent experiments with 3 to 5 mice/group/experiment. *P ≤ 0.05, **P ≤ 0.01, and ***P ≤ 0.001, by 2-tailed Student’s t test.

### Co-infection of SARS-CoV-2 and *Kp* induces severe inflammation in the lungs

Following the gross diagnosis of bronchointerstitial pneumonia as well as the recruitment of large numbers of inflammatory cells to the lungs (Fig. 4F), we performed histopathology on the lung at 6 dpi (2 days post-*Kp* infection). Bronchointerstitial pneumonia, thrombosis, fibrin, epithelial cell necrosis, perivascular cuffing, edema, and hemorrhage were the most common lung lesions (Fig. 5A-D). The lesions were graded on a scale of 0 to 4, with 0 being nonexistent and 4 being severe. Control and *Kp*-infected mice lungs had open alveoli and few histopathological changes (Fig. 5A and C). The lung infected with S-CoV-2 had multifocal, moderate interstitial pneumonia (black arrow, Fig. 5B) and perivascular cuffing (black arrowhead, Fig. 5B). The alveoli showed moderate edema and fibrin deposition. Conversely, the pathologic lesions in the co-infected mice were the most severe (Fig. 5D). The alveoli and bronchioles were densely packed with neutrophils and macrophages and were admixed with edema and fibrin (bronchopneumonia, black star, Fig. 5D). There was multifocal thrombosis in small caliber vessels. Innumerable bacteria were also observed in the lesions. Overall, S-CoV-2 + *Kp* mice showed significantly higher lesion scores than control, *Kp*, and S-CoV-2 (Fig. 5E).

**Figure 5.**
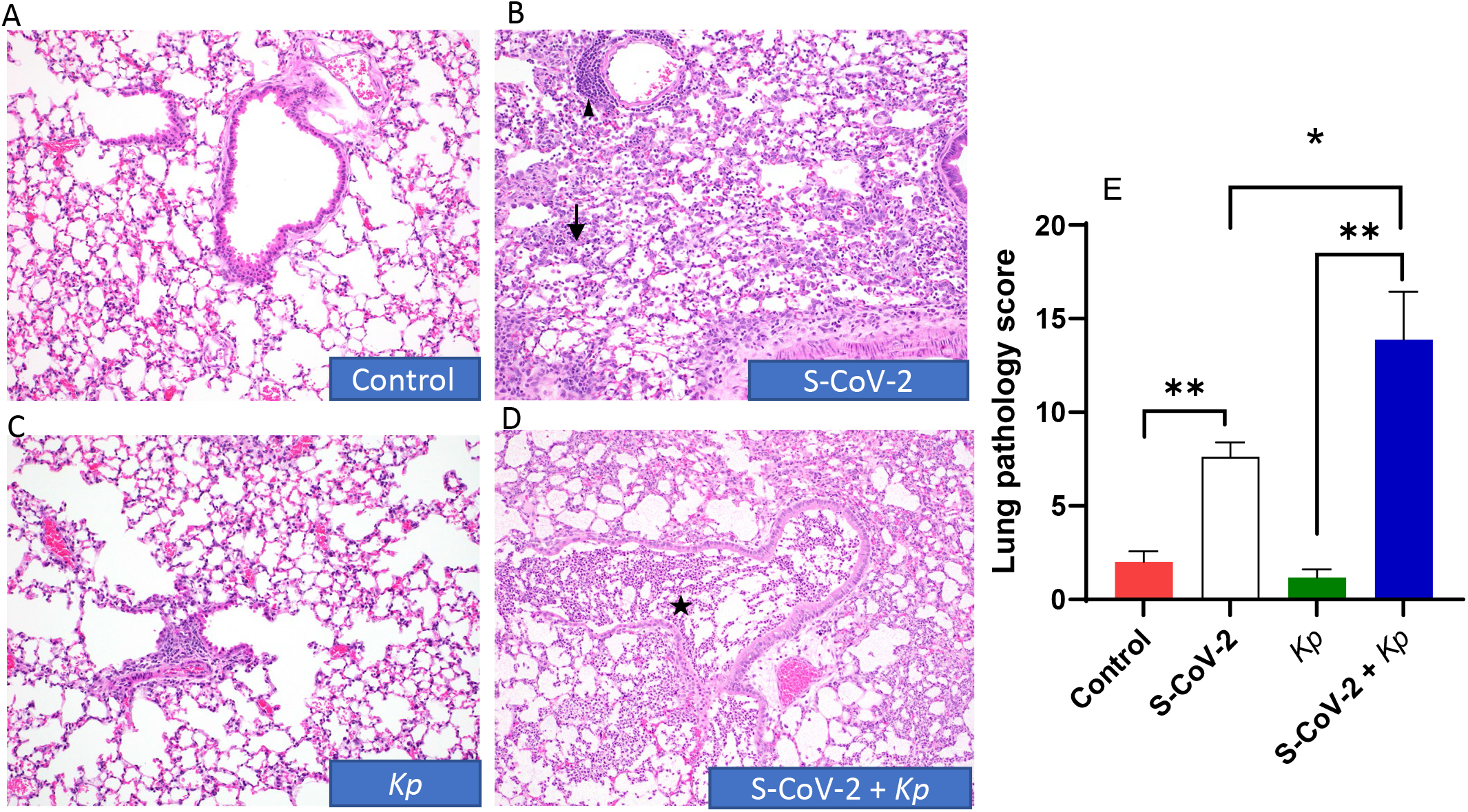
Lung pathology in co-infected mice. Photomicrographs of Lungs (A-D) at day 6 post-infection demonstrating lung pathology (Hematoxylin and Eosin stain) E. average of total lung pathology scores from two board-certified pathologists, n = 3-8 mice per group. P value < 0.05 based on a Welch’s test, S-CoV-2 vs S-CoV-2 + *Kp*, *Kp* vs S-CoV-2 + *Kp*, control vs S-CoV-2.

## Discussion

Respiratory co-infections pose a significant risk of exacerbating disease processes, primarily due to the overwhelming burden placed on the immune system by multiple pathogens. Each pathogen elicits a unique immune response, and the dysregulation that occurs in co-infections can impair the body’s ability to mount an effective defense against both viral and bacterial invaders. This further complicates accurate diagnosis and treatment [30], as the clinical manifestations of co-infections can be intricate and challenging to interpret. Therefore, gaining insight into the influence of co-infections on disease progression becomes crucial in guiding appropriate diagnostic and therapeutic approaches. Given the prevalence and severity of secondary bacterial infection during COVID-19, there is a growing need to understand the pathogenesis and impact of co-infections. In this study, we aimed to investigate the pathogenesis of co-infection with SARS-CoV-2 and *Kp* using a mouse model. We chose *Kp* for further coinfection studies based on its prevalence in multiple hospital reports [18, 19, 31] and our patient cohort (Fig. 1) as well as its potential for drug resistance [17]. We discovered that sublethal doses of *Kp* and SARS-CoV-2 can be lethal to mice when co-infected (SARS-CoV-2 followed by *Kp*). The mice exhibited severe clinical signs and weight loss along with increased mortality after 2-4 days of *Kp* infection. Histopathology demonstrated severe pneumonia along with a large influx of neutrophils, macrophages, and innumerable bacteria. Flow cytometric analysis also demonstrated a significant increase in the neutrophil and inflammatory monocytes/macrophage population.

In the case of COVID-19, coinfection with other respiratory viruses or bacteria can increase the risk of respiratory failure and death [16]. Furthermore, co-infection with SARS-CoV-2 and other viruses may also result in changes in the way the body responds to treatment. For example, studies have suggested that influenza virus co-infection may interfere with the effectiveness of SARS-CoV-2 antiviral medications [32]. Gram-positive bacteria such as *Streptococcus pneumoniae* (*Sp*) are commonly found in COVID-19 patients [33]. Similarly, gram-negative bacteria were detected in 64% of COVID-19 patients with pulmonary bacterial infection [26], which highlights their potential to cause secondary bacterial pneumonia. *Kp* is a gram-negative bacterium that can cause a range of infections, including pneumonia, urinary tract infections, and sepsis. It is often found in healthcare settings where it can be transmitted through contact with contaminated surfaces or medical equipment. Coinfection with both SARS-CoV-2 and *Kp* has been shown to cause severe illness [14, 18, 19, 31]. The emergence of drug-resistant Carbapenemase-producing *Kp* is a growing concern, with a high prevalence (34%) in intensive care unit patients in certain areas [18]. Additionally, there have been reports of patients hospitalized for SARS-CoV-2 infection who subsequently developed hypervirulent *Kp* strain infection, resulting in fatal outcomes [19, 34]. The global spread of antimicrobial resistance *Kp* [19, 35] and frequent reports of hospital-acquired infection [36, 37], underscores the need to study *Kp* pathogenesis in the context of COVID-19.

Mouse models of co-infection are key to understand the complex interactions between viruses, bacteria, and host innate and adaptive immunity. Several pathogens, such as influenza virus and respiratory syncytial virus [38], *Mycobacteria* [39], Human immunodeficiency virus [40], and *Streptococcus pneumoniae* [41], have been studied for co-infection with SARS-CoV-2. However, there are few animal models developed to understand the pathogenesis of SARS-CoV-2 and coinfections [23, 42–44]. Recent co-infection studies demonstrated increased susceptibility and lethality due to *Sp* and SARS-CoV-2 co-infection [23, 24]. In these studies, mice showed reduced survival rates due to increased bacterial loads in the lungs. Notably, mortality was observed regardless of whether the SARS-CoV-2 challenge occurred before or after *Sp* infection [23]. The increased bacterial levels after co-infection were associated with reduced alveolar macrophages, making co-infection more severe than either infection alone [23]. Neutrophil and macrophage/monocyte dysfunction have been reported in SARS-CoV-2 infections [45]. These cells are at the forefront of an antibacterial immune response; thus, dysfunctional phenotype in these cells predisposes COVID-19 patients to secondary bacterial infections. Neutrophils and monocytes isolated from critically ill SARS-CoV-2 patients failed to clear *Sp* and *Sa* [46], demonstrating impairment of antibacterial function [47]. These findings suggest that viral infection can impair antibacterial defense and enhance bacterial load, and this effect seems to be independent of whether the bacteria are gram-positive or gram-negative [23, 42] (Fig. 3A).

Respiratory viruses can trigger a cytokine storm, which can result in damage to the alveoli increasing leakage of fluid from the vessel [48] as well as systemic effects of inflammation and damage to other organs [49]. In severe cases of SARS-CoV-2 pneumonia, 42% patients had acute respiratory distress syndrome (ARDS), leading to hypoxia [50]. Individuals with severe COVID-19 demonstrate interstitial pneumonia with alveolar damage and hyaline membrane formation [51]. We observed rubbery firm lungs (gross pathology) with interstitial pneumonia, lung consolidation, infiltration of mononuclear cells, perivascular cuffing, and thickened alveolar septa in the lungs of SARS-CoV-2-infected mice. These lesions were consistent with other mouse models of SARS-CoV-2 infection [22, 52] indicating that our co-infection model largely mimicked the lung pathology associated with COVID-19. However, the lack of typical ARDS-like changes in the SARS-CoV-2-only group could be due to sublethal challenge as the morphological features of the disease vary based on the severity and duration of the disease. Nevertheless, in the case of co-infection with *Kp*, the pathological alterations became notably severe with the lungs exhibiting a substantial presence of inflammatory cells, along with an abundance of fibrin, numerous bacteria, and necrosis.

In summary, we have developed a mouse model of co-infection with SARS-CoV-2 and *Kp* and our study provides insights into the disease pathogenesis. The findings highlight the significance of monocyte and neutrophil dysfunction in secondary bacterial infections, leading to increased lung pathology and mortality. Strategies aimed at restoring the antibacterial activity of these cells may hold promise in preventing clinical complications associated with secondary bacterial infections in COVID-19 patients. Further research is needed to uncover the underlying mechanisms of COVID-19 severity following co-infection and develop targeted therapeutic approaches to mitigate the impact of secondary bacterial infections in COVID-19 patients.

## Acknowledgments

The following reagent was obtained through BEI Resources, NIAID, NIH: Cercopithecus aethiops Kidney Epithelial Cells Expressing Transmembrane Protease, Serine 2 and Human Angiotensin-Converting Enzyme 2 (Vero E6-TMPRSS2-T2A-ACE2), cat no. NR-54970. We also thank Dr. Dr. Stanley Perlman (University of Iowa) for providing Mouse-adapted SARS-CoV-2 strain. Research reported in this publication was supported by the National Institute Of General Medical Sciences of the National Institutes of Health under Award Number P20GM103648. The content is solely the responsibility of the authors and does not necessarily represent the official views of the National Institutes of Health. In addition, start-up funds from the College of Veterinary Medicine at Oklahoma State University were used for this study.

